# Cocaine-induced immediate-early gene expression in the nucleus accumbens: roles of separate cAMP sensors

**DOI:** 10.1101/2025.04.15.648980

**Authors:** Hai-Ying Zhang, Tabinda Salman, Guo-Hua Bi, Sunny Z. Jiang, Charles R. Gerfen, Andrew Lutas, Wenqin Xu, Zheng-Xiong Xi, Lee E. Eiden

**Author notes:** Correspondence to: Lee E. Eiden, Section on Molecular Neuroscience, NIMH Intramural Research Program, 9000 Rockville Pike, Building 49, Room 5A38, Bethesda, MD, USA, 20892, E-mail.

## Abstract

Immediate-early gene (IEG) induction after administration of amphetamine or cocaine has been used to trace the signaling pathways that mediate neuronal plasticity required for the short- and long-term behavioral effects of these psychostimulants. We recently reported that a novel cyclic AMP (cAMP)-dependent Rap guanine nucleotide exchange factor-2 (RapGEF2)-ERK signaling pathway is required for Egr-1 induction in D1 medium-spiny neurons (MSNs) of the nucleus accumbens (NAc) after cocaine treatment, and that its deletion from the NAc neurons attenuates cocaine-induced locomotor sensitization and conditioned place preference (CPP). However, the cell type-specific neuronal mechanisms underlying this effect remain unclear. In this study, we used Cre-LoxP technology and a novel Cre-amplifier transgene to generate conditional RapGEF2 knockout mice targeting D1-MSNs and investigated the functional role of RapGEF2 in cocaine reward. Deletion of RapGEF2 in D1-MSNs blocked cocaine-induced ERK phosphorylation (p-ERK) and Egr-1 induction. D1-MSN-specific RapGEF2 deletion did not affect intravenous cocaine self-administration, nor did it affect Fos induction by cocaine, prompting us to examine more closely the role of metabotropic (cAMP-dependent) signaling to IEGs after cocaine administration. We used a battery of D1-MSN-specific genetic interventions targeting cAMP signaling, including Drd1-Cre::Rap1A/B^fl/fl^ mice, and AAV injection of a Cre-dependent catalytically active phosphodiesterase (PDE4D3-cat) or a Cre-dependent protein kinase A-inhibitor (PKI) in the NAc of Drd1-Cre mice, to explore further the underlying cAMP dependence of IEG induction by acute and chronic cocaine administration, and the cAMP sensors required. Rap1 is reported as a necessary component for both RapGEF2- and PKA-dependent ERK activation, but a requirement for Rap in Fos induction by cocaine has not been examined. Deletion of Rap1A/B in D1-MSNs blocked cocaine-induced p-ERK and Egr-1 expression, but not c-Fos, supporting the idea that the RapGEF2-Rap1-ERK pathway specifically regulates Egr-1, not c-Fos, expression. D1-specific expression of the PDE4D3-cat ablated up-regulation of both Egr-1 and Fos in NAc after cocaine administration, demonstrating that induction of both IEGs requires cAMP elevation in D1-MSNs. Notably, inhibition of PKA activity via AAV-mediated expression of PKI-alpha in D1-MSNs blocked both c-Fos and Egr-1 induction. Thus, acute or chronic cocaine administration activates at least two cAMP-dependent signaling pathways in D1-MSNs: a PKA-Fos pathway and a RapGEF2-ERK-Egr-1 pathway. The finding that PKA also activates the ERK-Egr-1 signaling pathway by convergence on Rap1, and concomitantly activates c-Fos independently of Rap1, may underlie selective effects of metabotropic activation of RapGEF2 and PKA activation by cAMP on cocaine-dependent behaviors in mice.

## INTRODUCTION

Cocaine and amphetamine are psychostimulants that cause changes in brain state leading to compulsive drug seeking and taking behavior (addiction). The major action of psychostimulants in the brain is thought to be a dramatic increase in extracellular dopamine in the striatum. In fact, most of the effects of these stimulants have been conceptualized as arising from by short- and long-term changes in the neurochemistry of dopamine D1 and D2 receptor-expressing medium spiny neurons (MSNs) of the NAc, which is richly innervated by mesencephalic dopaminergic projections. D1-MSNs have been consider the predominant targets involved in the development of cocaine addiction (Lu *et al*. 2006), and this is consistent with attenuation of the rewarding effects of psychostimulants in D1 receptor-deficient mice (Caine *et al*. 2007), and interference with Drd1 mRNA translation directly in the NAc (Pisanu *et al*. 2015). A prominent effect of response to cocaine exposure is immediate-early gene (IEG) up-regulation, predominantly in D1-MSNs, in the NAc (Lu *et al*. 2006; Pickens *et al*. 2011). This intracellular response to increased DA and D1 receptor stimulation is likely to be important in mediating the cellular plasticity important for development of drug addiction.

The pathways through which IEGs are up-regulated after psychostimulants have been studied intensively to provide potential mechanisms for cellular plasticity underlying short- and long-term effects including locomotor sensitization, and increased place preference associate with drug ingestion (Lu *et al*. 2006; Valjent *et al*. 2000; Valjent *et al*. 2006b; Valjent *et al*. 2006a; Xu 2008; Zhang *et al*. 2006). Egr-1 (also called Zif268 and NGFI-A) has emerged as an IEG whose induction is postulated to be causal in psychostimulant action, and several studies have focused on effects of Egr-1 on cocaine-induced behaviors (Chandra & Lobo 2017; Valjent *et al*. 2006a). Fos is also robustly induced in the NAc after psychostimulant administration (Graybiel *et al*. 1990; Drago *et al*. 1996). The role(s) of individual IEGs in mediating cocaine-induced late gene regulation important in psychostimulant long-term action, however, have not yet been determined (Savell *et al*. 2020).

D2-MSNs of the NAc are also appear to contribute to the rewarding effects of cocaine (Self 2014; Lobo & Nestler 2011; Heinsbroek *et al*. 2017; Gong *et al*. 2021; Chen *et al*. 2022; Mews *et al*. 2024), reflecting the importance of D1-D2 interactions the overall striatal response to psychostimulant exposure (Lobo & Nestler 2011). To better understand how D1- and D2-MSNs act, and interact, in mediating psychostimulant actions and behaviors, more precise delineation of the signaling cascades activated by cocaine, and specific to each neuronal, is required. This means elucidation of the linkages between the release of first messengers, the elevation of second messengers, the induction of IEGs, the expression of late genes, and the changes in neuronal activity patterns, that finally result in the behaviors associated with psychostimulant intake and drug-seeking, in a cell-type specific way. This in turn, requires methodology that distinguishes between D1-mediated effects of dopamine directly at D1-MSNs, and those that are mediated indirectly, through D2-dependent intercellular signaling to D1 neurons, following psychostimulant administration.

Copious literature has demonstrated that both calcium and cAMP are able to induce IEG expression in neurons in response to various stimuli (Morgan & Curran 1995; Sheng & Greenberg 1990; Hiroi *et al*. 1999). In response to cocaine, Fos and Egr-1 induction have been variously ascribed to activation of Rap1 via RasGRP1 (Nagai *et al*. 2016a; Nagai *et al*. 2016b), RapGEF2 (Jiang *et al*. 2021), and DARP-32 (Svenningsson *et al*. 2004; Hiroi *et al*. 1999; Zachariou *et al*. 2006; Zachariou *et al*. 2002). However, the molecular linkages between dopamine-dependent D1 receptor activation, cAMP, cAMP sensors, and individual IEGs, comprising the signaling cascades for induction of each one, are incompletely defined. Thus, it is critical to examine the mechanisms of metabotropic signaling in D1-MSNs in the NAc using genetic manipulations that are specific to this neuronal subpopulation. Here, we developed mouse strains in which RapGEF2 or Rap1A/B are ablated in D1-MSNs of striatum. We also generated mice in which cAMP elevation or PKA activation was specifically disrupted in D1-MSNs. Using these transgenic mice, we examined IEG expression following acute or chronic cocaine treatment as a function of elimination of various signaling components downstream of cAMP. IEGs such as Egr-1 and Fos can be induced by first messengers including GPCR ligands, excitatory neurotransmitters like glutamate, or neurotrophins such as BNDF (Morgan & Curran 1986; Sheng *et al*. 1988; Sheng & Greenberg 1990; Greenberg *et al*. 1992; Sheng *et al*. 1993; Robertson *et al*. 1995; Segal & Greenberg 1996; Zhao *et al*. 2007; Inaba *et al*. 2023). We tested whether Fos induction after treatment with cocaine, like Egr-1 induction (Jiang *et al*. 2017; Jiang *et al*. 2021), is due to metabotropic cAMP-dependent signaling, and, if so, what cAMP sensor mediates Fos induction. We used D1-MSN-specific RapGEF2 knockout to determine if development of chronic drug self-administration, leading to robust drug preference, was dependent upon this cAMP-dependent signaling pathway for ERK activation and Egr-1 induction.

## RESULTS

Both metabotropic and ionotropic first messengers can activate signaling pathways in D1-MSNs that result in the induction of immediate early genes (IEGs), including c-Fos and Egr-1 after psychostimulant administration. Locomotor response and the corresponding IEG induction profiles for psychostimulant administration for 1-6 days in C57Bl6/J mice are shown in Supplementary Figure 1A. Consistent with previous findings, locomotor activity is increased by either cocaine or methamphetamine on first i.p. injection, and increased further by subsequent injections over 5 days, of either drug (locomotor sensitization; Supplementary Figure 1B). Fos and Egr-1, as well as other members of the Fos and Egr gene families, were induced by psychostimulants both acutely (first injection, Coc1 or Meth1) and after six days of daily injection (Coc6 or Meth6), including both Egr-1 and −2, and JunB and c-Fos in the Fos-related gene family. None of these IEGs exhibited desensitization of induction upon repeated cocaine administration (Bcl-2 shown as a reference gene, Supplementary Figure 1C).

It has been previously shown that deletion of RapGEF2 in the NAc using AAV-mediated expression of Cre recombinase in RapGEF2^fl/fl^ mice, blocked ERK phosphorylation and Egr-1 induction, and eliminated cocaine-induced behaviors including locomotor sensitization and CPP (see Figure 1A, and (Jiang *et al*. 2021)). We wished to examine the effects of RapGEF2 deletion only from D1-MNS on cocaine-induced IEG expression, and behaviors. To this end, RapGEF2^fl/fl^ and Drd1-Cre mouse lines were crossed with the intention to create mice in which RapGEF2 was deleted from D1-expressing neurons. TaqMan and *in situ* hybridization probes (Supplementary Figure 2A) targeting exon 4 of RapGEF2 were designed to allow detection of loss of exon 4 after its Cre-dependent excision in the NAc in Drd1-Cre::RapGEF2^fl/fl^ mice.

**Figure 1.**
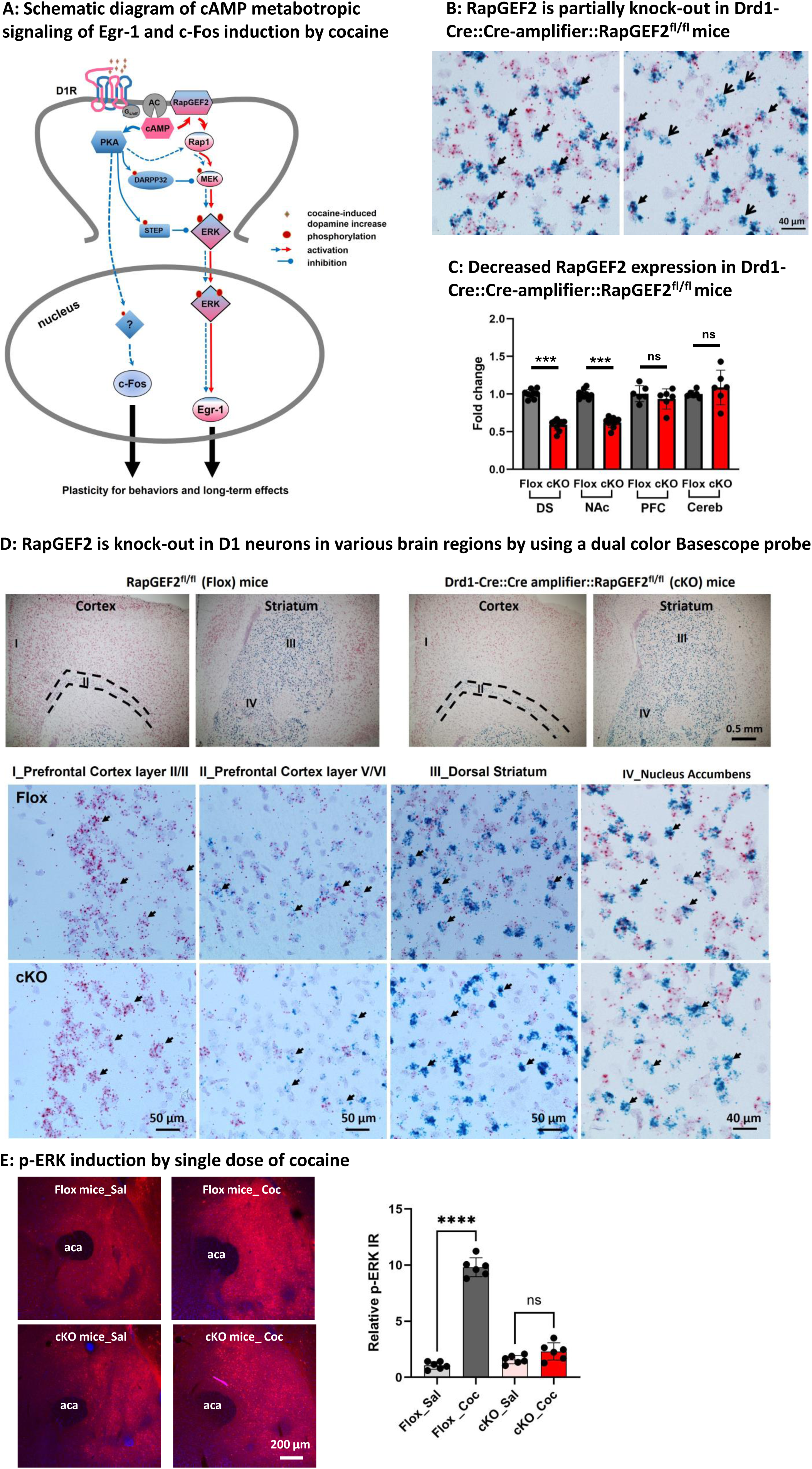
Generation of D1 specific RapGEF2 knock-out mice and their responses to cocaine. (A) Schematic diagram of metabotropic signaling pathway in D1 MSNs for the Egr-1 and c-Fos induction by cocaine. Cocaine acts by increasing extracellular dopamine leading to two major metabotropic signaling pathway: D1→cAMP→PKA (in Blue) and D1→cAMP→RapGEF2→Rap1→ERK (in Red), to modulate IEG, c-Fos and Egr-1 expression, resulting in the plasticity for behaviors or long-term effects. (B) qRT-PCR data showed about 40% decrease of RapGEF2 mRNA expression in both dorsal striatum (DS) and nucleus accumbens (NAc), but not in prefrontal cortex (PFC) and cerebellum (Cereb) as the control tissue in Drd1-Cre::CGW::RapGEF2^fl/fl^ mice, compared to Flox control mice. N = 5∼8 mice for each group. ***p < 0.001, ***p < 0.0001 for cKO versus Flox mice. All panels: Mean ± S.E.D. (C) Detection of RapGEF2 mRNA (shown by red color) in D1-MSNs (shown by blue color) in cortical and striatal areas. RapGEF2 mRNA are unbiquiously expressed in prefrontal cortex, dorsal striatum and ventral striatum both in RapGEF2 flox and Drd1-Cre::CGW::RapGEF2^fl/fl^ (cKO) mice shown in the low magnificent field. I-IV are the representative images from prefrontal cortex II/III (I), prefrontal cortex V/VI (II), dorsal striatum (III) and nucelues accumbens (IV). There is no RapGEF2 expression difference in prefrontal cortex layer II/III between flox and cKO mice (shown by solid arrow in I). While, in D1 neurons present brain regions, we did see the colocalization of RapGEF2 with Drd1a in flox control mice, but only see the Drd1a expressing D1 neurons without red dots (RapGEF2 mRNA) or non-D1 cells with the red dots (RapGEF2 mRNA) in cKO mice in II-IV, indicating the RapGEF2 was only knock-out in D1 neurons, but not other cells. (D) Representative images of p-ERK immunostaining after acute cocaine administration (15min after injection) both in flox control mice and Drd1-Cre::Cre-amplifier::RapGEF2^fl/fl^ (cKO) mice. Quantification data showed that a dramatic increase of p-ERK was present only in flox mice, but not in cKO mice. A total of 3∼4 sections from Bregma 1.6 mm to 1.2 mm of each mouse were included for quantification (n=3∼4 mice in each group). ****p < 0.0001 for cKO versus Flox mice. n.s: no statistic difference. All panels: Mean ± S.E.D.

Unexpectedly, the expression of intact RapGEF2 mRNA persisted in D1 neurons of Drd1-Cre::RapGEF2^fl/fl^ mice (Supplementary Figure 2B and Figure 1B), even though there was efficient excision of RapGEF2 from exon 4, and deletion of RapGEF2 mRNA, in excitatory neurons in CamK2α-Cre::RapGEF2^fl/fl^ mice (see (Jiang *et al*. 2021) and Supplementary Figure 2B), Both qRT-PCR probe and Basescope ISH probes showed a significant decrease of RapGEF2 both in cortex and hippocampus CA1 and DG area, but not in CA2 and CA3 area (Supplementary Figures 2B and C), as previously reported using RapGEF2 immunohistochemistry and Western immunoblotting (Jiang *et al*. 2024). However, RapGEF2 expression persisted in striatum of Drd1-Cre::RapGEF^fl/fl^ mice, indicating lack of Cre recombinase penetration to RapGEF2 loxP sites in Drd1-Cre::RapGEF2^fl/fl^ mice, and the need for a different strategy for RapGEF2 knockout in D1 neurons.

### Generation of D1-specific RapGEF2 knock out mice

To effectively knockout RapGEF2 in D1 neurons, we designed a Cre-dependent Cre-amplifier construct, Syn-Flox-STOP-Cre-Venus. This construct consists in a floxed STOP cassette, flanked by two loxP sites, inserted into a Cre coding sequence, downstream of a synapsin promotor (Supplementary Figure 3A). We bred Cre-amplifier^+^ mice with Drd1-Cre^+^ mice to generate a Drd1-Cre::Cre-amplifier strain, in which excision of the STOP cassette allows additional Cre expression under a strong synapsin promoter (Supplementary Figures 3B and C). Venus expression (detected with a GFP antibody) was robust in the dorsal and ventral striatum and sparsely distributed in the cortex, bed nucleus of the stria terminalis (BNST), hippocampus and paraventricular nucleus (PVN), confirming D1-Cre-specific expression of Cre from the Cre-amplifier gene. Drd1-Cre::Cre-amplifier::RapGEF2^fl/fl^ mice were generated by breeding Drd1-Cre::RapGEF2^fl/fl^ with Cre-amplifier::RapGEF2^fl/fl^ mice (Supplementary Figure 3D).

RapGEF2 expression was deleted from D1-expressing neurons in the Drd1-Cre::Cre-amplifier::RapGEF2^fl/fl^ mice. As shown in Figure 1C, there was a ∼40% decrease in RapGEF2 mRNA in the dorsal striatum (DS) and nucleus accumbens (NAc), consistent with the proportion of Drd1-compared to non-Drd1-expression neurons in striatum, but no detectable decrease in prefrontal cortex (PFC) and cerebellum (Cereb) where the percentage of Drd1^+^ cells is low and non-detectable, respectively (Gerfen *et al*. 2013). Dual color ISH demonstrated that RapGEF2 mRNA was only deleted from D1 neurons (labeled with drd1a, blue color) in the striatum and prefrontal cortex (Figure 1D). p-ERK induction by a single dose of cocaine was measured in the rostral NAc, where psychostimulant administration strongly affects ERK activation (Supplementary Figure 4). Deletion of RapGEF2 in D1 neurons dramatically attenuated p-ERK induction, in the rostral NAc after a single dose of cocaine (Figure 1E).

### Cocaine-induced Egr-1, but not c-Fos, expression is RapGEF2-dependent

The effect of RapGEF2 deletion from D1-MSNs on IEG induction after cocaine administration is shown in Figure 2. Based on the time course of Egr-1 and c-Fos induction after either acute or repeated cocaine treatment (Figures 2A and B), Egr-1 and c-Fos immunoreactivity (ir) was measured at 60 and 30 min, respectively, after cocaine administration. As expected, Egr-1 induction is abrogated by elimination of RapGEF2 from D1-MSNs. However, RapGEF2 deletion failed to alter acute or chronic cocaine-induced Fos induction (Figures 2B and C), consistent with previous observations that CREB phosphorylation, also a presumptive downstream target of PKA in D1-MSNs after cocaine administration, is unaffected by loss of RapGEF2 in the NAc (Jiang *et al*. 2021).

**Figure 2.**
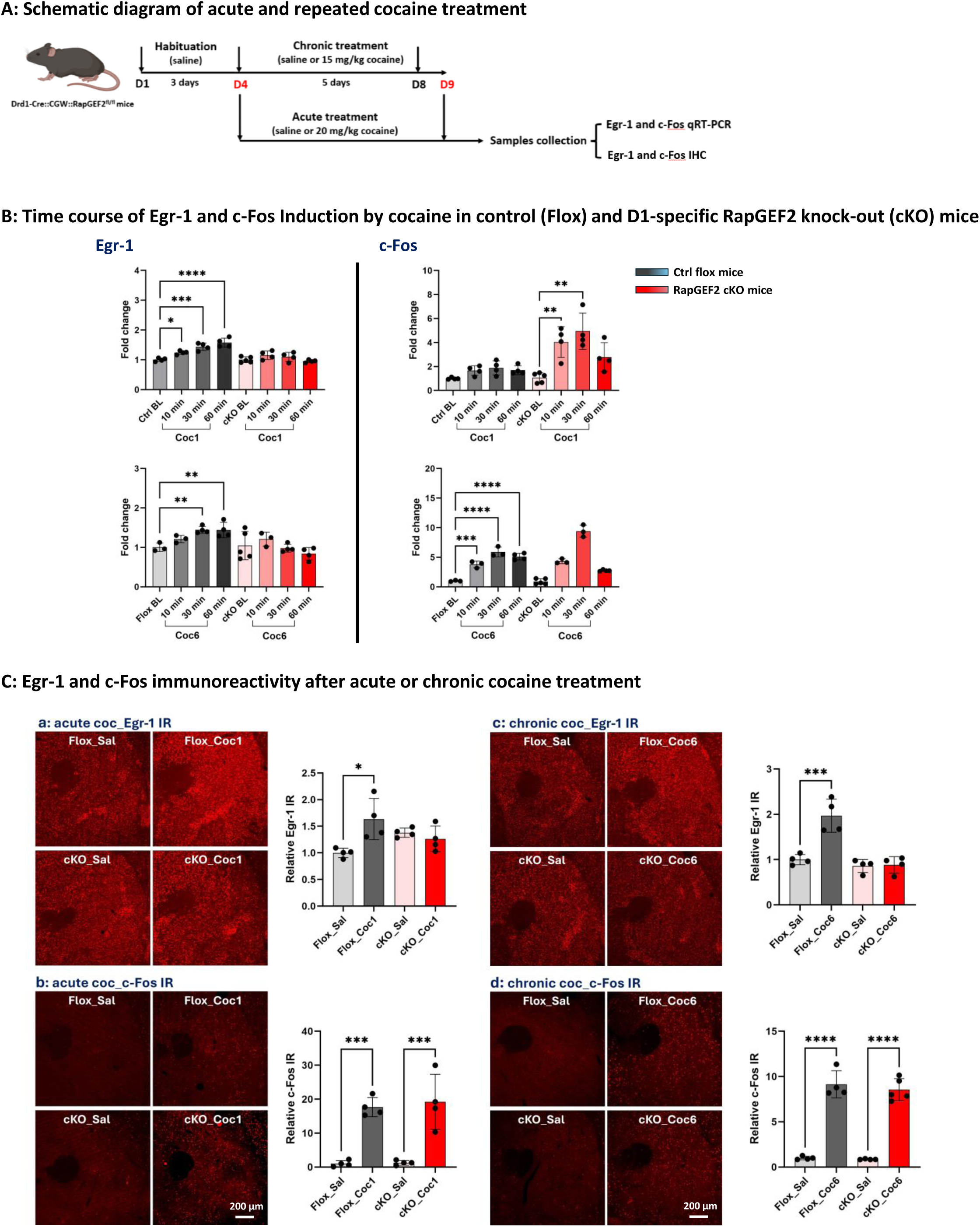
Induction of Egr-1, but not c-Fos, by cocaine is RapGEF2-dependent. (A) Schematic diagram of acute (20 mg/kg, i.p) and repeated cocaine (15 mg/kg, i.p) treatment for 6 consecutive days, tissue was collected at 10min, 30min or 60min after last injection of cocaine for c-Fos and Egr-1 detection either by qRT-PCR or immunohistochemistry assay. (B) Time course of Egr-1 and c-Fos induction by single dose of cocaine (Coc1) and 6 injections of cocaine (Coc6) in Flox control (Ctrl) and D1-specific RapGEF2 knock-out (cKO) mice at baseline (BL), 10min, 30min and 60min time point after last injection of cocaine. Gray color for Ctrl group and red color for cKO group. The peak induction of Egr-1 is at 60min no matter it’s the first injection or 6 injections of cocaine in Ctrl mice and cKO mice didn’t show the Egr-1 injection at all (Left). While the peak induction of c-Fos is at 30 min both in Ctrl mice and cKO mice (Right). N = 4∼5 mice for each group. *p < 0.05, **p < 0.01, ***p < 0.001, ****p < 0.0001 for different time point versus BL. All panels: Mean ± S.E.D. (C) Representative images and relative immunoreactivity (IR) quantification of Egr-1 (Top) at 60 min and c-Fos ir (Bottom) at 30 min after single dose of cocaine injection (Left) and last cocaine injection (Right). A total of 3∼4 sections from Bregma −1.2mm to 0.8 mm of each mouse were included for quantification (n=4 mice in each group). *p < 0.05, ***p < 0.001, ****p < 0.0001 for cocaine versus saline. All panels: Mean ± S.E.D.

### Deletion of RapGEF2 from D1-MSNs did not alter cocaine self-administration

Several laboratories have shown that development of place preference for cocaine is an ERK-dependent process, correlated with and likely dependent upon, induction of Egr-1 (Valjent *et al*. 2006b; Lu *et al*. 2006; Ferguson *et al*. 2006; Fasano *et al*. 2009; Fricks-Gleason & Marshall 2011; Philibin *et al*. 2011; Pickens *et al*. 2011; Kim *et al*. 2011; Jiang *et al*. 2021). We subjected RapGEF2^fl/fl^ (control flox) and D1-Cre::Cre-amplifier::RapGEF2^fl/fl^ (cKO) mice to a protocol in which they were allowed to self-administer cocaine intravenously. As shown in Figure 3, wild-type mice exhibited acquisition and maintenance of cocaine self-administration as assessed by both fixed-ratio (FR) and progressive-ratio (PR) reinforcement schedule, demonstrated by active and inactive lever response, cocaine infusions, and break-point level. Acquisition and maintenace of cocaine self-administration behavior was unimpaired in D1-specific RapGEF2 cKO mice.

**Figure 3.**
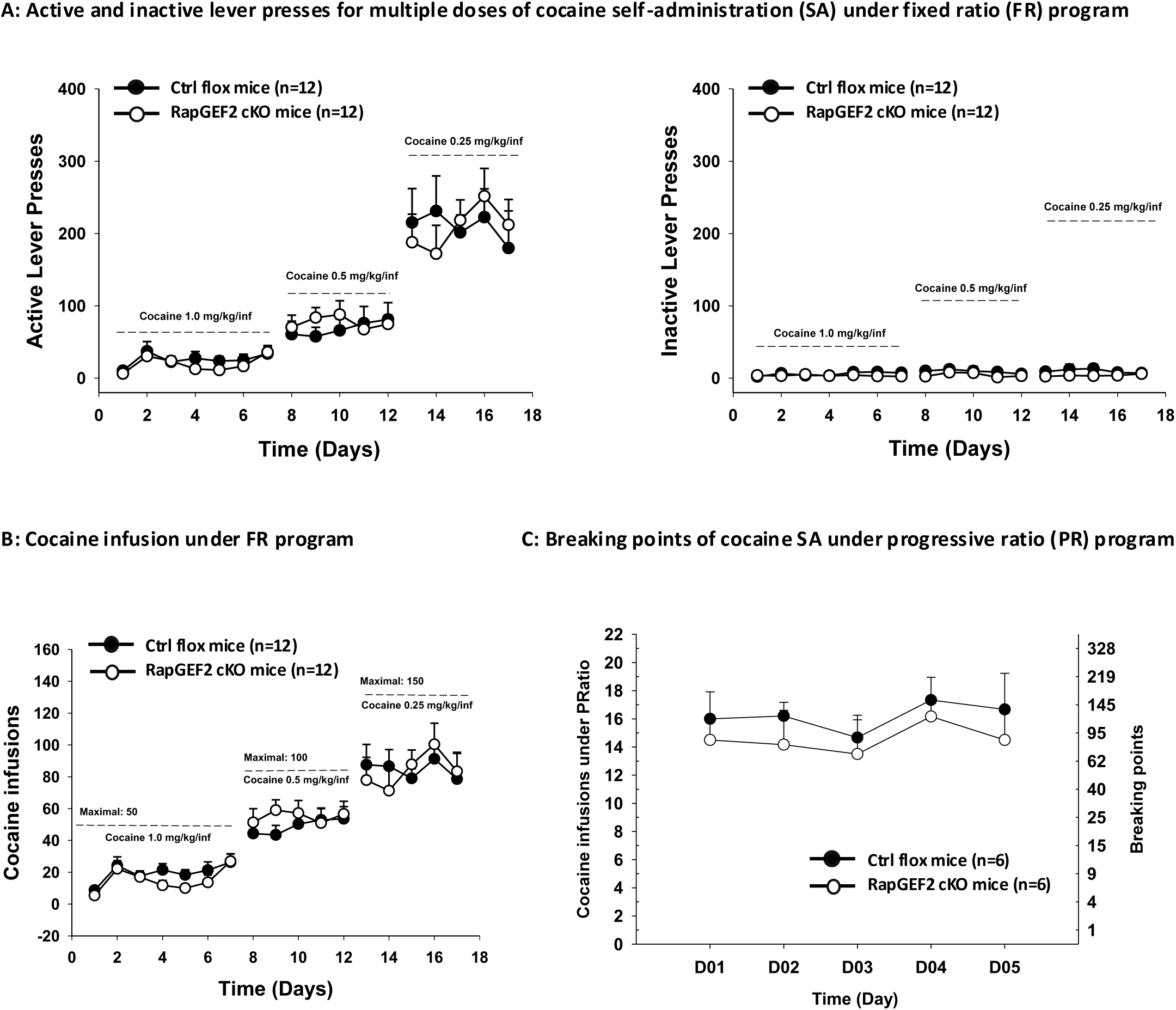
Cocaine taking and seeking behavior was not impaired in D1-specific RapGEF2 KO mice. (A) Cocaine-taking behavior in Flox control (Ctrl flox) mice and Drd1-Cre::CGW::RapGEF2^fl/fl^ (cKO) mice during acquisition of cocaine self-administration (SA) by using multiple dose of cocaine under fix ratio (FR) program. Active (Left) and inactive (Right) lever responding in each phase of the experiment, demonstrating a cocaine-dose-orderly increase of lever presses both in Ctrl and cKO mice and all the mice developed cocaine SA behavior. (B) Total number of cocaine infusions in daily test sessions during the acquisition phase of cocaine self-administration. (C) Cocaine self-administration under progressive ratio (PR) reinforcement, illustrating both Ctrl and cKO mice have similar motivation for cocaine shown by a total cocaine infusion and breaking points.

### Cocaine-induced up-regulation of p-ERK and Egr-1, but not Fos, is Rap1A/B-dependent

It has been proposed that PKA activation results in a Rap1-dependent activation of the MEK-ERK pathway (see Figure 1A; (Nagai *et al*. 2016a; Nagai *et al*. 2016b)). As a number of laboratories have postulated an ERK-dependent pathway for activation of Fos through phosphorylation of CREB (Grewal *et al*. 2000; Saito *et al*. 2013; Li *et al*. 2016), the dependence for Fos induction by cocaine on abrogation of Rap1A/B function was next examined. Crossing of mice in which both alleles of both Rap1A and Rap1B are floxed (Pan *et al*. 2008) with Drd1-Cre mice results in an about 50% decrease in Rap1 in the NAc assessed by Western blot assay (Figures 4A and B), consistent with loss of expression in D1-MSNs, and Rap1 A and B mRNA expression is ablated specifically in D1-MSNs in the NAc and throughout the brain, assessed by ISH (Figure 4C). Deletion of Rap1A/B expression in D1 neurons resulted in loss of ERK activation and Egr-1 induction, but with preservation of induction of Fos, in NAc, after acute administration of cocaine (Figure 4D).

**Figure 4.**
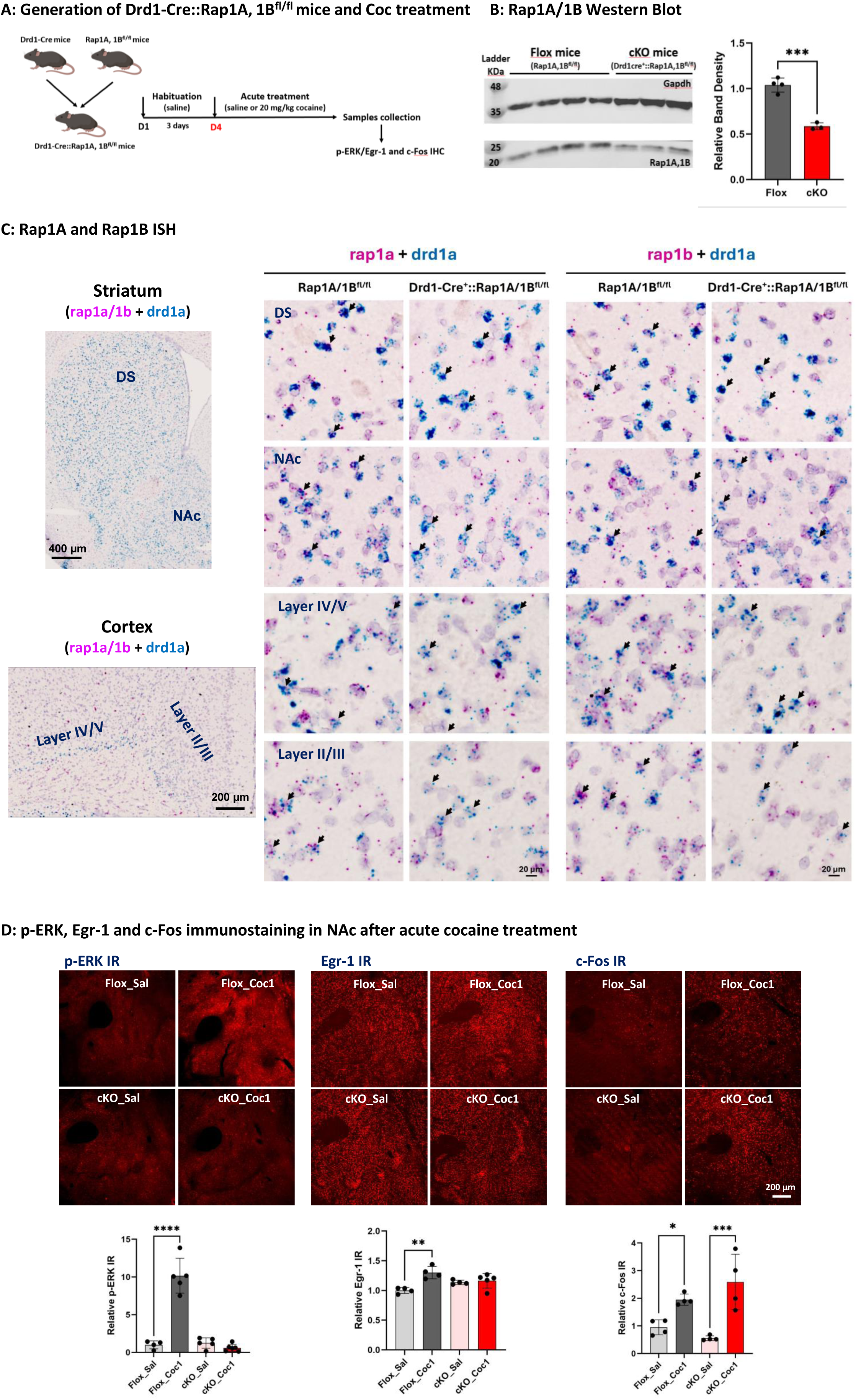
p-ERK, Egr-1, but not c-Fos, induction by cocaine is also D1 Rap1A/B-dependent. (A) Demonstration of Drd1-Cre::Rap1A/B^fl/fl^ mice generation by breeding Drd1-Cre mice with Rap1A/B^fl/fl^ mice and acute cocaine (20 mg/kg, i.p) treatment for p-ERK, Egr-1 and c-Fos detection. (B) Rap1A/B Western Blot showing around 40% decrease of Rap1A/B expression in NAc of Drd1-Cre::Rap1A/B^fl/fl^ (cKO) mice, compared to Rap1A/B^fl/fl^ (Flox) mice. n=3∼4 mice in each group, ***p < 0.001 for cKO versus ctrl mice. (C) Basescope ISH demonstrating that low to moderate level of Rap1a (Left, red dots) and Rap1b (Right, red dots) are expressed in both non-D1 dopamine neurons and D1 dopamine neurons (labeled with drd1a, blue dots) in dorsal striatum (DS), nucleus accumben (NAc) and cortex deep layer V/VI and layer III/IV, where the D1 dopamine neurons are present in flox mice. In the cKO mice, no either Rap1a or Rap1b was detected in drd1a labeling D1-neurons (shown by blue color). While, the Rap1a or Rap1b was still seen in non-D1 cells (with red dots only). The representative neurons in each group are shown by black arrows. (D) Induction of p-ERK, Egr-1, but not c-Fos are impaired in cKO mice after acute cocaine treatment. 10∼15 min for p-ERK, 30 min for c-Fos and 60 min for Egr-1 were used for tissue collection. A total of 3∼4 sections from Bregma 1.7 mm to 1.2 mm of each mouse were included for quantification (n=3∼4 mice in each group). *p < 0.05, **p < 0.01, ****p < 0.0001 for cocaine versus saline. All panels: Mean ± S.E.D.

### Cyclic AMP dependence for both Fos and Egr-1 induction by cocaine administration

Egr-1 induction after psychostimulant administration requires cAMP-dependent signaling components (Nagai *et al*. 2016a; Jiang *et al*. 2017). To determine whether Fos induction after cocaine treatment is due to ionotropic (non-cAMP-dependent) versus metabotropic (cAMP-dependent) signaling, we examined the effects of suppression of cAMP action in D1-MSNs on Fos elevation after cocaine administration. A constitutively active form of the catalytic domain of phosphodiesterase 4 (PDE4D3-cat) was expressed using a Cre-dependent AAV vector, unilaterally in the NAc, and induction of Fos and Egr-1 protein in the NAc was measured 30 min after cocaine injection, and compared to the corresponding area on the uninjected side (Figure 5A). Both Fos and Egr-1 induction after cocaine requires cAMP elevation, as evidenced by suppression of its induction in cells expressing PDE4D3-cat (Figure 5B).

**Figure 5.**
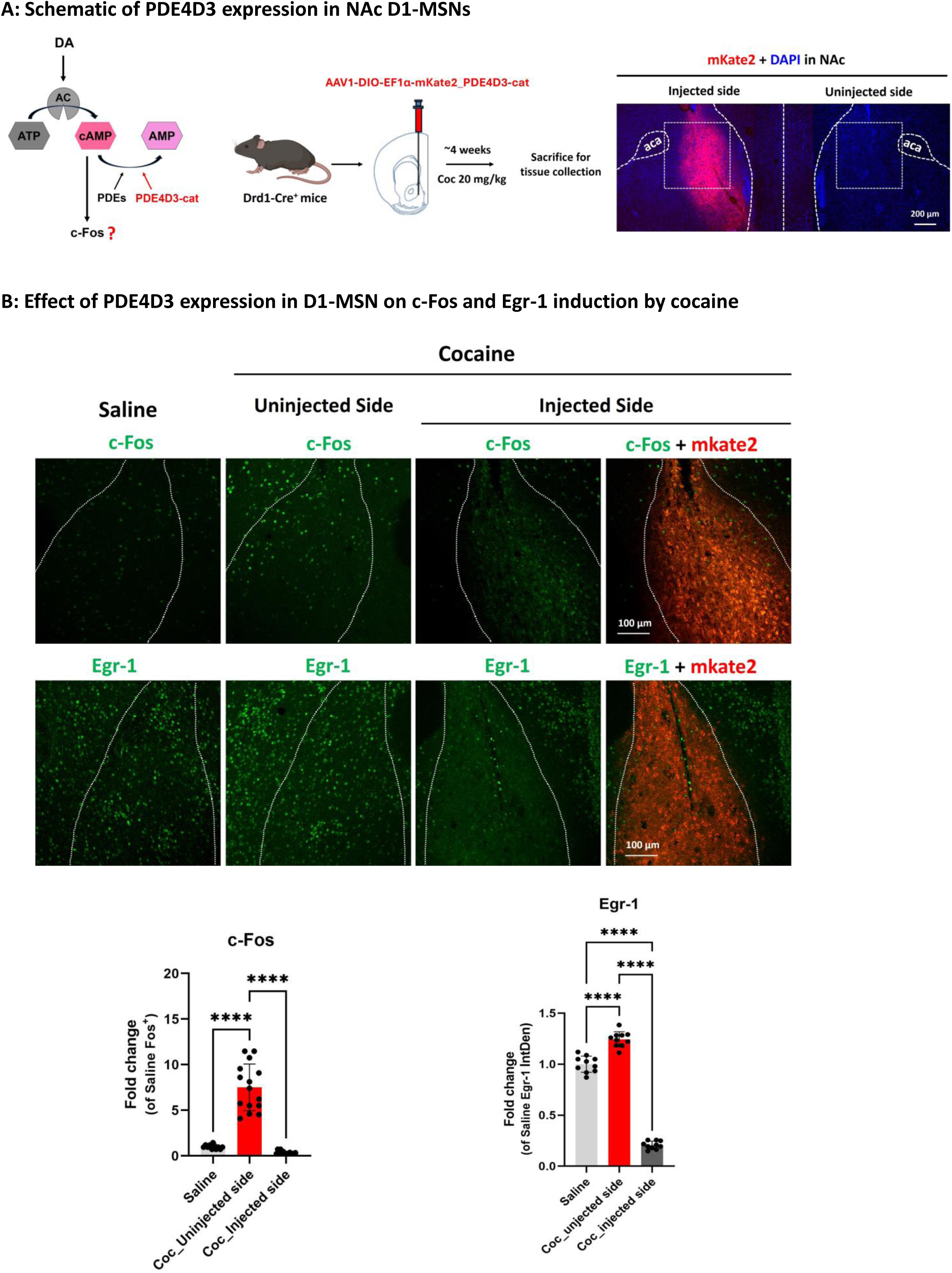
Breakdown of cAMP production blocks both Egr-1 and c-Fos induction by cocaine. (A) Schematic of phosphodiesterase 4D3 catalytic domain (PDE4D3-cat) enhancing the breakdown of cAMP production following cocaine administration (Left) and Drd1-Cre^+^ mice receiving unilateral PDE4D3-cat injections (shown by mKate2 red epifluoresence) were challenged with cocaine injection for c-Fos and Egr-1 detection (Right). (E) Representative images for c-Fos-ir (Top, green) and Egr-1-ir (bottom, green) with or without mkate2 epifluorescence (Red) in both injected side and uninjected side after saline or cocaine administration. Integrated density of c-Fos or Egr-1 was measured in unilateral NAc of saline groups and cocaine groups with or without PDE4D3-cat virus injection. A total of 3∼4 unilateral sections from Bregma −1.2mm to 0.8 mm of each mouse were included for quantification (n=3 mice, a total of 10∼15 fields in each group). ****p < 0.0001 for Coc versus saline or uninjected side versus injected side. All panels: Mean ± S.E.D.

### PKA dependence of IEG induction after cocaine

Induction of Fos and Egr-1 after cocaine both require elevation of cAMP within D1-MSNs. We further investigated whether PKA activation was required for IEG induction after cocaine, via expression of the PKA inhibitor PKIα in D1-MSNs of ventral striatum (Figure 6A). As expression of PKIα (as evidenced by visualization of mRuby from the PKIα-mRuby bicistronic message) is stringently Cre-dependent, we crossed Drd1-Cre mice with mice expressing the Cre-amplifier (Supplementary figures 3B and C). Since the Cre-amplifier gene allows Venus expression specifically in D1-expressing neurons, Drd1-Cre::Cre-amplifier mice allowed demonstration that PKIα, after injection of AAV-Flex-PKIα-IRES-nls-mRuby2 (Chen *et al*. 2014) into NAc, was expressed only in D1-expressing cells (Figure 6B). In these mice, IEG expression in D1-MSNs not expressing PKIα and those expressing PKIα (indicated by mRuby) could be compared after treatment with cocaine. Inhibition of PKA by overexpression of the endogenous protein kinase A inhibitor PKIα in D1-MSNs of NAc resulted in attenuation of both Fos and Egr-1 induction after cocaine (Figure 6C). These results show that while Egr-1 induction (Jiang *et al*. 2021) requires the cAMP sensor RapGEF2, the latter is necessary but may not be sufficient on its own for cAMP-dependent Egr-1 induction, and that PKA also plays a role in this signaling pathway.

**Figure 6.**
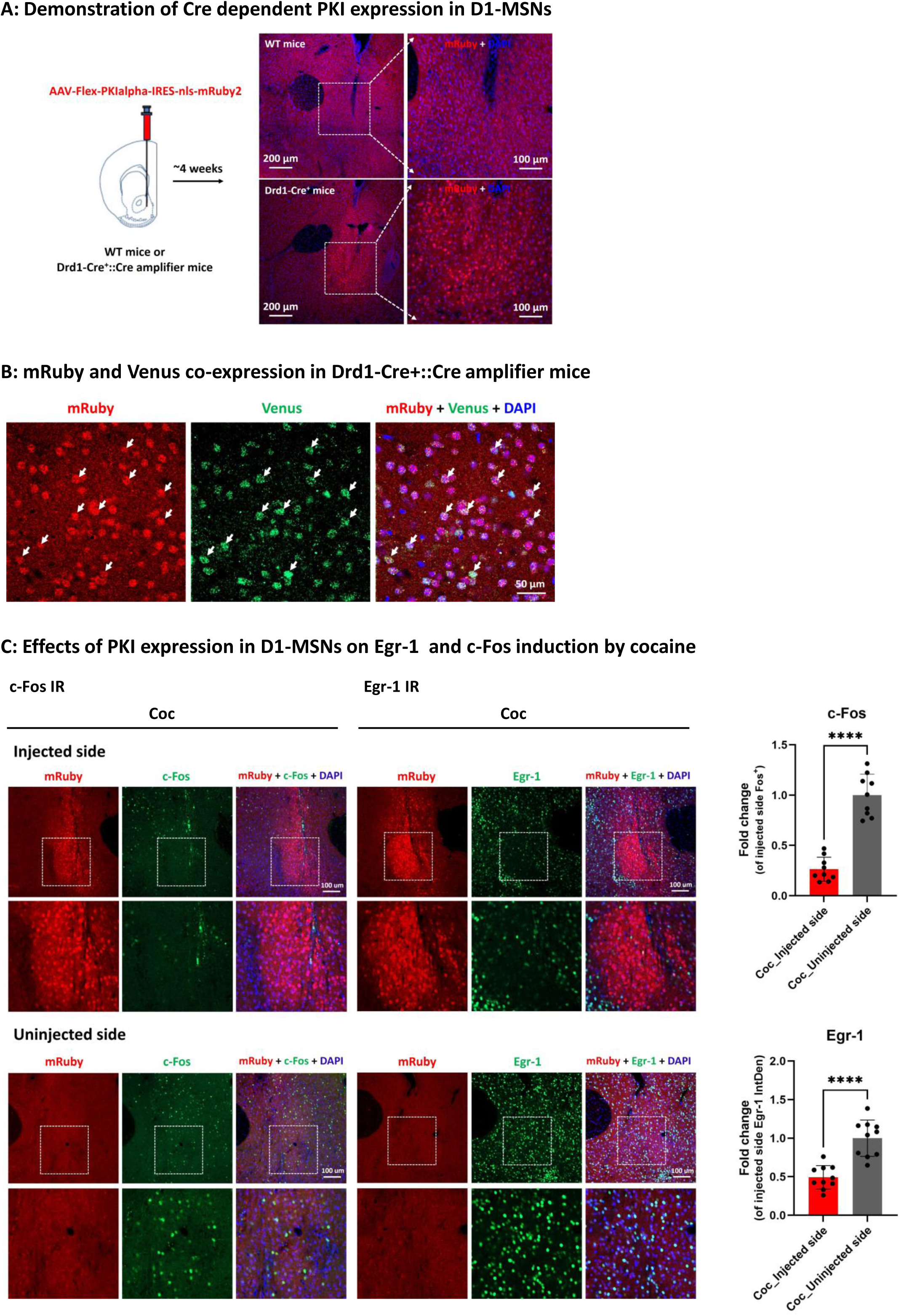
Effects of inhibition of PKA signaling on Egr-1 and c-Fos induction by cocaine. (A) WT or Drd1-Cre^+^ or Drd1-Cre^+^::Cre amplifier mice receiving unilateral injection of Flex-PKI-mRuby virus into NAc and only Cre^+^ mouse showed mRuby expression. (B) Drd1-Cre^+^::Cre amplifier mice receiving unilateral injection of Flex-PKI-mRuby virus into NAc showed the colocalization of mRuby and Venus by IHC (shown by white arrow), indicating PKI was selectively expressed in D1-MSNs in a Cre-dependent manner. (C) Representative images of c-Fos (Right) or Egr-1 (Left) induction by acute cocaine (20 mg/kg, i.p) administration after Drd1-Cre^+^ mice received unilateral PKI virus injection into the NAc. Fold change of integrated density (IntDen) of c-Fos or Egr-1 was measured in unilateral NAc section of cocaine groups with or without PKI virus injection. A total of 3∼4 unilateral sections from Bregma 1.7 mm to 1.3 mm of each mouse were included for quantification (n=3 mice, a total of 10 fields in each group). ****p < 0.0001 for injected side versus uninjected side. All panels: Mean ± S.E.D.

## DISCUSSION

In this report, we have used a variety of genetic tools to modulate cAMP levels, and the activities of two downstream cAMP sensors, PKA and RapGEF2, to examine the relationship between metabotropic activation of D1-MSNs, IEG induction, and behavioral consequences after administration of cocaine. We have identified differential regulation of the IEGs Egr-1 and Fos on the basis that cAMP-dependent induction of Egr-1 is Rap1-dependent, while Fos induction is Rap1-independent. Thus, D1-MSN-specific deletion of RapGEF2 results in differential impairment of cocaine-dependent Egr-1 induction without affecting that of Fos. The same pattern is observed after D1-MSN-specific deletion of Rap1A/B. Suppression of cAMP elevation after treatment with cocaine results in impairment in induction of both IEGs, as does inhibition of PKA. Delineation of this pattern of IEG activation through cAMP activated signaling may help to illuminate the causal roles of specific IEGs in late gene activation required for cellular plasticity underlying behavioral consequences of long-term cocaine administration or self-administration.

Cocaine-induced hyperactivity and self-administration of cocaine (Caine *et al*. 2007) are abolished or attenuated in Drd1-deficient mice (Xu *et al*. 1994; Abraham *et al*. 2016). The requirement for D1 receptor activation for the neuronal plasticity underlying both long- and short-term effects of cocaine and related psychostimulants, is also supported by the effects of D1-specific antagonists (Schindler & Carmona 2002; Bertran-Gonzalez *et al*. 2008). IEG induction in NAc, specifically in D1-MSNs, is a hallmark of both D1 agonist treatment, and cocaine administration (Teague & Nestler 2022). Since both Fos and Egr-1 are implicated in behavioral effects of cocaine, attention to the precise mechanisms whereby Fos and Egr-1 are up-regulated by psychostimulant administration has received a great deal of attention (Gao *et al*. 2017; Teague & Nestler 2022; Xu 2008). The regulation of both IEGs is complex, and has been postulated to require first messenger signaling not only by dopamine but by glutamate (Keefe & Gerfen 1996; Valjent *et al*. 2005). With this in mind, we have examined in detail the downstream effects of cAMP elevation in D1-MSNs of NAc, to identify the IEG signatures of metabotropic signaling associated with cocaine administration.

We have previously shown that ERK phosphorylation and Egr-1 induction after cocaine requires the cAMP sensor RapGEF2, while CREB phosphorylation after cocaine administration does not require RapGEF2 (Jiang *et al*. 2021). However, it has not been explicitly demonstrated that cAMP elevation per se is required for RapGEF2 activation, nor whether cocaine-dependent Fos and Egr-1 induction require the same or distinct cAMP sensors. In fact, there are conflicting reports on the role of cAMP sensors and the downstream effectors involved in Fos induction after cocaine exposure. Maximal induction of Fos by acute administration of cocaine at 20 mg/kg is reported to be unaffected in mice bearing a mutated DARPP-32 incapable of activation by PKA (Zachariou *et al*. 2006). On the other hand, cocaine induction of deltaFosB induction does require expression of DARPP-32 (Hiroi *et al*. 1999), and amphetamine induction of Fos in the dorsal striatum is blocked by CREB antisense RNA, suggesting a cAMP-dependent signaling pathway for Fos activation by this psychostimulant (Konradi *et al*. 1994). Here we have created mice in which cAMP elevation can be suppressed by expression of PDE4D3, a catalytically active cAMP phosphodiesterase (Singh Alvarado *et al*. 2024), in D1 neurons, in order to determine whether both Egr-1 and Fos induction by cocaine require cAMP. Furthermore, PKA is required for activation of both IEGs, as evidenced by abrogation of their induction in NAc D1-MSNs expressing the PKA inhibitor PKIα. Deletion of Rap1, a previously postulated intermediate in ERK activation by PKA after activation of D1 receptors, resulted in inhibition of Egr-1 induction by cocaine, but spared Fos elevation, confirming dual activation of Egr-1 induction by cAMP through both PKA and RapGEF2. It is noteworthy that activation of Fos by cocaine, however, is completely Rap1 independent.

Finally, we tested the ability of mice with RapGEF2 permanently deleted from D1 neurons to successfully develop stable cocaineself-administration behavior. RapGEF2-deleted mice displayed a similar pattern as wild-type mice in the acquisition and maintenance of cocaine self-administration. This result was surprising, as it has been postulated that Egr-1 is critical for cocaine-induced CPP in mice (Valjent *et al*. 2006a), and that RapGEF2 deletion from the NAc after AAV-Cre injection of RapGEF2^fl/fl^ mice abrogates CPP as well (Jiang *et al*. 2021). There are several possibilities for our findings. One is that while deletion of RapGEF2 from the NAc eliminates Egr-1 induction in D1-MSNs, this alone is insufficient to eliminate behavioral effects of cocaine administration, either passively or self-administered, unless RapGEF2 is absent from both D1- and D2-MSNs (Jiang *et al*. 2021). A second possibility is that cocaine self-administration leading to acquisition of cocaine preference depends upon a brain state, e.g. heightened sensitivity to reward reinforcement during foraging behavior, habit formation, etc., that does not exist during passive cocaine administration (Kalivas & Volkow 2005; Everitt & Robbins 2005; Zapata *et al*. 2010). This process may lead to altered behavior in a way that is more heavily dependent on PKA-than on RapGEF2-dependent activation of Egr-1. A third possibility is that elimination of RapGEF2 early in development, versus in adult mice, allows a compensatory pathway for cocaine reward registration in D1-MSNs that is independent of RapGEF2, and perhaps independent of Egr-1 induction altogether, requiring instead induction of Fos or other RapGEF2-independent IEGs. In any case, it may be that Fos activation is dominant over that of Egr-1 in eliciting gene transcription mediating cocaine reinforcement during self-administration, compared to registration of drug preference after repeated passive cocaine administration. If so, it will be worthwhile to investigate, using newly developed tools for manipulation of cAMP levels in the ventral striatum, whether metabotropic, rather than ionotropic induction of either Fos or Egr-1 (Morgan & Curran 1986; Sheng & Greenberg 1990; Greenberg *et al*. 1992; Sheng *et al*. 1993; Robertson *et al*. 1995) represents the dominant first-messenger input to D1-MSNs for mediating cocaine-taking behavior. The investigations reported here illuminate mechanisms of D1-mediated IEG activation after psychostimulant administration, and suggest additional investigations to probe cAMP-dependent IEG induction in D1-MSNs as a potential key to psychostimulant drug-taking and seeking behaviors.

## MATERIALS AND METHODS

### Animals and Drugs

Male or female mice (WT or transgenic) on C57BL/6J background were housed in a maximal of 5 per cage and acclimatized to 12 h light/12 h dark cycle with food and water ad libitum. Animal cares were approved by the National Institute of Mental Health Institutional Animal Care and Use Committee and conducted in accordance with the National Institutes of Health guidelines. The Drd1a BAC-Cre driver line (Drd1-Cre^+/−^-FK150, MGI:3836633), the floxed RapGEF2 (RapGEF^fl/fl^) mouse strain, Drd1-Cre::RapGEF2^fl/fl^ and corticolimbic RapGEF2 conditional knock-out (cKO) mouse strain Camk2α-cre^+/−^::RapGEF2^fl/fl^ were generated as described previously (Gerfen *et al*. 2013; Jiang *et al*. 2021; Jiang *et al*. 2017). Rap1A/B double floxed (Rap1A/B^fl/fl^) mice were generated as described previously (Pan *et al*. 2008) and crossed with Drd1-Cre mice to produce Drd1-Cre^+^::Rap1A/B^fl/fl^ mice by a Cre-LoxP system (Figure 4A). Pharmaceutical grade cocaine, methamphetamine and D1 agonist SKF81297 were purchased from Millipore Sigma. Cocaine and methamphetamine are directly dissolved in saline for a working solution. SKF81297 was first dissolved in DMSO to get 4 mg/ml stock solution and stored at −20℃ freezer, then diluted in saline for a working solution before injection. All drugs used *in vivo* were administrated by intraperitoneal injection, except for the intravenous cocaine self-administration.

### RapGEF2 Conditional Knock-out (cKO) Mouse Generation

#### Cre-amplifier plasmid construct and mouse line generation

We designed a Cre dependent Cre amplifier construct, also call Syn-Flox-STOP-Cre-Venus, in which a floxed STOP cassette, flanked by two loxP sites, was inserted into a Cre coding sequence downstream of a synapsin promotor, and inserted randomly into the genome (Supplementary Figure 3A). In the founder line, a total of 4 primers sets was used to identify the presence of 4 major components, including Synapsin promoter, Cre sequence, STOP cassette and Venus sequence of the construct and DNA sequencing was further used to confirm the correction of the whole construct sequence. We bred Cre^+^ amplifier mouse with Drd1-Cre^+^ mouse to generate Drd1-Cre:Cre amplifier mouse, in which the STOP cassette was excised to allow more Cre expression under a strong Synapsin promoter, instead of Drd1a promotor and the D1 neurons were also lighted up with Venus expression (Supplementary Figures 3B and C), by using a GFP antibody.

#### D1-specific RapGEF2 cKO mice

Cre amplifier or CreGoneWild (CGW)::RapGEF2^fl/fl^ mice were generated by breeding RapGEF2^fl/fl^ with CGW^+^ mice. All off springs need to be genotyped both for Cre and Flox to choose CGW^+^::RapGEF2^fl/fl^, to further bred with RapGEF2 Floxed homozygous mice. Finally, the Drd1-Cre^+^::RapGEF2^fl/fl^ and CGW^+^::RapGEF2^fl/fl^ mice were used to set up breeders (Supplementary Figure 3D). The off springs, both flox homozygous, as the control mice and Drd1-Cre^+^::CGW^+^::RapGEF2^fl/fl^, as the cKO mice, were used for experiment. All transgenic mouse lines primers are listed in Table 1

**Table 1:**
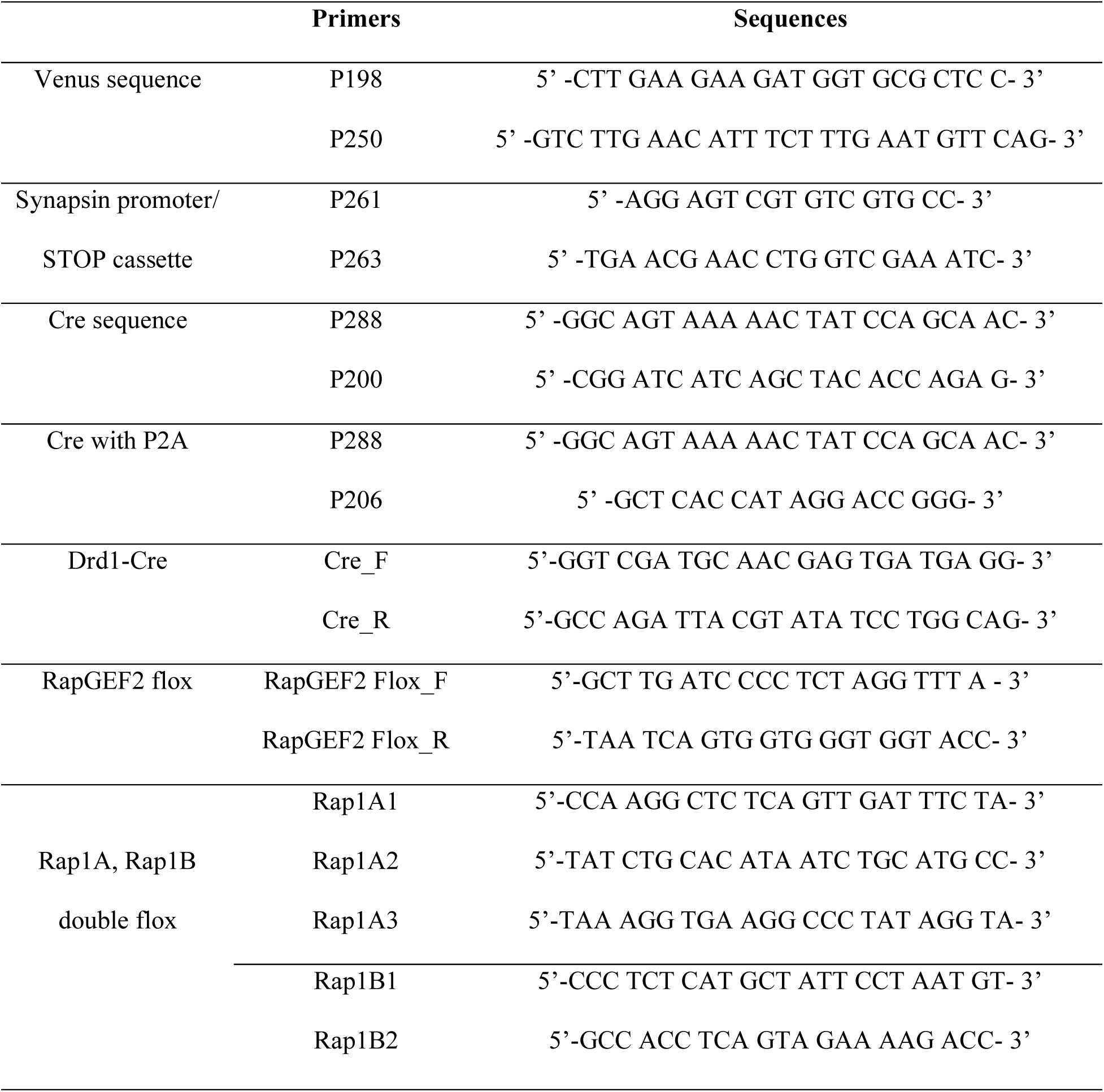
Primers for transgenic mice.

### Viral Injection

AAV9.CAG.Flex.tdTomato.WPRE.bGH control virus was obtained from Penn Vector Core. AAV1-Flex-PKIalpha-IRES-nls-mRuby2 was purchased from Addgene (Plasmid #63059) and packaged by VigeneBiosciences. AAV1-DIO-EF1α-mKate2-PDE4D3-cat was kindly provided by Dr. Andrew Lutas in National Institute of Diabetes and Digestive and Kidney Diseases (NIDDK), NIH. Drd1-Cre^+^ or Drd1-Cre^+^::Cre^+^ amplifier mice were unilaterally injected with either tdTomato control virus, under the control of the CAG promoter or PKIalpha-mRuby2 virus, expressing PKIalpha and nuclear localized mRuby2, to inhibit PKA signaling pathway or mKate-PDE4D3-cat virus, expressing PDE4D3-cat to break down the cAMP, in the NAc in a Cre-dependent manner. Surgeries and viral injection were conducted according to the “NIH-ARAC guidelines for survival rodent surgery”. Briefly, animals were anesthetized, shaved and mounted into the stereotaxic apparatus. A small midline incision was then made and the pericranial tissue was teased away from the skull with an ethanol-soaked swab to enable identification of the Bregma and Lambda areas. Using a small hand-held drill, a very small hole was made in the skull according to the calculated coordinates for NAc (Bregma 1.32 mm, ML 0.7 mm, DV 4.7mm). A Hamilton syringe (pre-loaded with viral particles) was then slowly lowered, penetrating the dura to the determined depth. A volume of 0.4 μl of the virus (∼1.0×10^12^ infectious particle per microliter) was slowly (∼0.08 μl/min) injected into the brain. A 5-minute wait is performed before the needle is very slowly retracted from the brain to prevent backflow of the viral vector. The animals were allowed to recover and subjected to drug treatment four weeks after viral injection. Brain injection sites were histologically verified with counterstaining from post-mortem sections and plotted on standardized sections from the stereotaxic atlas.

### Immunohistochemistry and Microscopy

Immunohistochemistry was conducted as previously described (Jiang *et al*. 2021) after animal perfusion with freshly made 4% paraformaldehyde. Briefly, mouse brains were sectioned by a Vibratome at a 40 μm thickness. Free-floating sections were washed 3 times in the wash buffer, either PBS containing 0.3% Triton X-100 (PBST) or TBS containing 0.5% Triton X-100 (TBST) (15 min for each), incubated at room temperature in blocking solution (10% normal goat or donkey serum in wash buffer; 2 h), and then incubated with primary antibodies in blocking solution overnight at 4 ℃. The following day, sections were washed 3 times in wash buffer (15 min for each), incubated in the dark in Alexa 488 or 555-conjugated secondary antibodies for 2 h. Sections were mounted in Vectashield (Vector Laboratories, Burlingame, CA). Confocal images were obtained on a Zeiss LSM 510 confocal microscope at the National Institute of Neurological Disorders and Stroke Light Imaging Facility. The primary antibodies used were rabbit anti-c-Fos (EnCor Biotechnology Inc., Gainesville, FL, Cat# RPCA-c-Fos, 1:5,000), rabbit anti-Egr-1 (Cell Signaling Technology, Inc, Danvers, MA, Cat# 4154, 1:1,000), rabbit anti-p-ERK (Cell Signaling Technology, Danvers, MA, Cat# 4370, 1:1,500), rabbit anti-GFP (Invitrogen, Waltham, MA, Cat# A-11122, 1:500), goat anti-mRuby (Genprice Inc. San Jose, CA, Cat#AB3410, 1:500), goat anti-tdTomato (MyBiosource, Inc, San Diego, CA, Cat# MBS448092, 1:1,000), Alexa 488- or 555-conjugated goat anti-rabbit or donkey anti-goat IgG (Millipore Sigma, Burlington, MA, 1:500)

### Brain Dissection

Mice were sacrificed by a quick cervical dislocation, and whole brains were quickly removed from skull. After a brief rinse in the ice-cold PBS, the brains were placed into a pre-cold mouse brain matrix in 500 µm thick slices and to be sectioned. The hippocampal CA1 was dissected out from slice from approximately Bregma − 1.43 mm to bregma − 2.53 mm by a scalpel and forceps under dissecting microscope. A tissue block from striatum area was first obtained from approximately Bregma 1.8 mm to bregma − 0.8 mm by using a brain matrix. The bilateral dorsal striatum (DS) and ventral striatum (NAc) were punched out by using 1 mm punch needle. Other brain area, including whole hippocampus, prefrontal cortex (PFC) and cerebellum (Cereb), were also collected (∼20mg) as reference area. Structure boundaries were identified using a mouse brain atlas (Franklin & Paxinos 2019). Once dissected, samples were immediately snap-frozen in Dry Ice and later stored at – 80℃ until further qRT-PCR or Western Blot assay.

### Western Blot

The NAc punched samples were sonicated on ice with RIPA buffer supplemented with Halt Protease and Phosphatase Inhibitor Cocktail and centrifuged at 3,000 rpm for 10 min at 4°C. Supernatants were collected, and protein concentration was determined by MicroBCA Protein Assay kit (Thermo Fisher Scientific) according to the manufacturer protocol. Western blot was performed as described previously (Jiang *et al*. 2021). The rabbit anti-Rap1A/B (Upstate, catalog #; 1:500) and anti-GAPDH (D16H11) (Cell Signaling Technology, catalog #5174; 1:1,000) antibodies were used for the blotting. IR bands were visualized with a SuperSignal West Dura Chemiluminescent Substrate (Thermo Fisher Scientific), photographed with a ChemiDoc Imaging system, and quantified with National Institutes of Health ImageJ.

### RNA extraction, cDNA synthesis, and qRT-PCR

Total RNA was extracted using the RNeasy Mini kit (cat no. 74104, Qiagen, Germantown, MD, USA) following the manufacturer’s instructions. RNA concentration was evaluated by absorbance at 260 nm using the NanoVue Plus spectrophotometer (Biochrom US, Holliston, MA, USA). RNA samples were used for cDNA production when A260/A280 ratios were between 1.8 and 2.1 and stored at −80°C until use. Single-stranded cDNAs were synthesized with SuperScript III First-Strand Synthesis SuperMix (cat no. 11752-059, Thermo Fisher Scientific, Waltham, MA, USA), according to the kit protocol, using 0.5 μg total RNA. TaqMan gene expression assay for RapGEF2 (Mm01219742_m1) and endogenous Gapdh (catalog # 4352661) were purchased from Thermo Fisher Scientific. qRT-PCR protocols were as reported previously (Zhang *et al*. 2014; Jiang *et al*. 2021). Briefly, the qRT-PCR reactions were set up in triplicate and performed in a Bio-Rad iCycler system (cat no.170-8720, Bio-Rad, Hercules, California, USA) using the following PCR program: 95°C hold for 20 s; 40 cycles of 95°C; denaturation for 3 s, and 60°C annealing and extension for 30 s. Quantification used the reference gene Gapdh with the 2^−ΔΔCt^ method. Primers are listed in Table 2

**Table 2:** TaqMan probes for qRT-PCR.

| Gene Symbol | Assay ID | Cat# |
| --- | --- | --- |
| Gapdh | Mm99999915_g1 | 4448489 |
| Rapgef2 | Mm01219742_m1 | 4351372 |
| Dusp1 | Mm05854205_s1 | 4351372 |
| Fos | Mm00487426_g1 | 4331182 |
| Bcl2 | Mm00477631_m1 | 4331182 |
| Nr4a3 | Mm00450071_g1 | 4331182 |
| egr-2 | Mm00456650_m1 | 4331182 |
| egr-1 | Mm00656724_m1 | 4331182 |
| Junb | Mm01251660_s1 | 4331182 |

### Basescope Chromogenic in situ Hybridization (ISH)

*In situ* hybridization was conducted using the RNAScope 2.5 HD Duplex Detection Kit (Chromogenic, cat no. 323,100, ACD Bio, Newark, CA, USA), as directed in the manufactory manual. Briefly, fresh-frozen mouse brains were sectioned coronally on a cryostat to 12 µm. Sections were mounted on Superfrost Plus slides (Fisher Scientific, Waltham, MA, USA). Slides were fixed with freshly made 4% paraformaldehyde for 30 min at 4 ℃, then dehydrated in an ethanol gradient. Hydrogen peroxide and protease treatments, probe hybridization and signal development proceeded as described in the manual. Custom designed BA-Mm-Drd1a-2zz-st-C1 (Cat No. 1097761-C1), BA-Mm-RapGEF2-2zz-st-C2 (cat No. 1098111-C2), BA-Mm-Rap1A-2zz-st-C2, BA-Mm-Rap1B-2zz-st-C2 probes were developed as blue for drd1a and red color for rapGEF2, rap1A and rap1B, respectively. Images were obtained on a bright-field microscope.

### Behavioral Tests

Locomotor activity was conducted in NIMH Rodent Behavioral Core Facility. Cocaine intravenous self-administration tests was conducted in NIDA IRP.

#### Locomotor activity

WT mice were tested for locomotor activity in responses to single and multiple doses of cocaine and methamphetamine. Briefly, mice were placed in an Accuscan open field chamber (20 × 20 cm^2^, 8 beam sensors for each dimension) to monitor locomotor activity. Room light was set up as 30-50 lux. For the first 4 days, mice were placed for 30 min in the activity chamber (habituation phase), received an injection of saline (i.p.), then placed back to the chamber for 1 h. Then locomotor activity induced by psychostimulants was tested by replacing injection of saline with cocaine (15 mg/kg, i.p.) or methamphetamine (2 mg/kg, i.p.) for 5 consecutive days in the same setting.

#### Intravenous (i.v.) cocaine self-administration (SA)

##### Surgery

RapGEF2^fl/fl^ (Ctrl flox) and Drd1-Cre^+^::CGW^+^::RapGEF2^fl/fl^ (cKO) mice were prepared by jugular vein catheterization for i.v. cocaine self-administration as described previously. Briefly, catheterization was performed under sodium pentobarbital anesthesia using standard aseptic surgical techniques and the self-administration cannulae were fixed to the skull with 4 stainless steel jeweler’s screws (Small Parts, Miami Lakes, FL) and dental acrylic cement. During experimental sessions, each catheter was connected to an injection pump via tubing encased in a protective metal spring from the head-mounted connector to the top of the experimental chamber. To help prevent clogging, the catheters were flushed daily with a gentamicin– heparin saline solution (0.1 mg/ml gentamicin and 30 IU/ml heparin; ICN Biochemicals, Cleveland, OH).

##### Multiple doses of cocaine SA under fix ratio (FR) reinforcement

After 2 weeks recovery from surgery, mouse was placed into a test chamber (daytime-dark phase) and allowed to lever-press for i.v. cocaine (1 mg/kg/infusion) delivered in 0.01 ml over 4.6 s on a fixed ratio 1 (FR1) reinforcement schedule. Each cocaine infusion was associated with presentation of a stimulus light and tone. During the 4.6 s infusion time, additional responses on the active lever were recorded but did not lead to additional infusions. Each session lasted 3 h. FR1 reinforcement was used for 5 days until stable cocaine self-administration was established as described previously (Song *et al*. 2012; Zhang *et al*. 2014). Mice were then allowed to continue cocaine self-administration under FR2 reinforcement with first 0.5 mg/kg/infusion, then 0.25 mg/kg/infusion, each for 5 days. To avoid cocaine overdose, each animal was limited to a maximum of 50 cocaine injections for FR1 and 100 cocaine injections for FR2 during 3 h session.

##### Cocaine SA under progressive ratio (PR) reinforcement

After stable cocaine self-administration under FR2 reinforcement was established, the mice were switched to cocaine (0.5 mg/kg/injection) self-administration under a PR schedule, during which the work requirement (lever presses) needed to receive a single i.v. cocaine infusion was progressively raised within each test session (Zhang *et al*. 2014; Song *et al*. 2012) according to the following PR series: 1, 2, 4, 6, 9, 12, 15, 20, 25, 32, 40, 50, 62, 77, 95, 118, 145, 178, 219, 268, 328, 402, 492, and 603 until breakpoint (BP) was reached. BP was defined as the maximal workload (lever presses) completed for a cocaine infusion before a 1-h period during which no infusion was obtained by the animal. The PR schedule is computer programmed to progress to a maximum of 603 and the average BP was ∼150. Animals were allowed to continue daily sessions of cocaine self-administration under PR reinforcement conditions for 5 consecutive days.

### Statistical analysis

Mice of both female and male were used in all studies. The sample size (n) per group is described in the figure legends for each experiment. Statistical analyses were conducted using GraphPad Prism 10. Student’s t tests and one-way or two-way ANOVAs were used where appropriate. Post hoc analyses were performed using the Bonferroni test. Data were reported using histograms and scatter plots to represent mean ± S.E.D. and individual data in each group. Differences were significant when p < 0.05.

## STATEMENTS & DECLARATIONS

## Author Contributions

All authors contributed to the study conception and design. Material preparation, data collection and analysis were performed by Hai-Ying Zhang, Tabinda Salman, Guo-Hua Bi, Wenqin Xu, and Sunny Jiang. All authors wrote and commented on the previous versions of manuscript, and read and approved the final manuscript.

## Data Availability

The datasets generated and analyzed during the current study are available from the corresponding author on reasonable request.

## Ethics approval

Animal studies was approved by the NIMH Institutional Animal Care and Use Committee (IACUC) and conducted in accordance with NIH guidelines.

## Consent to participate

No human subjects were involved.

## Consent to publish

No human subjects were involved.

---

**Supplementary Figure 1.**
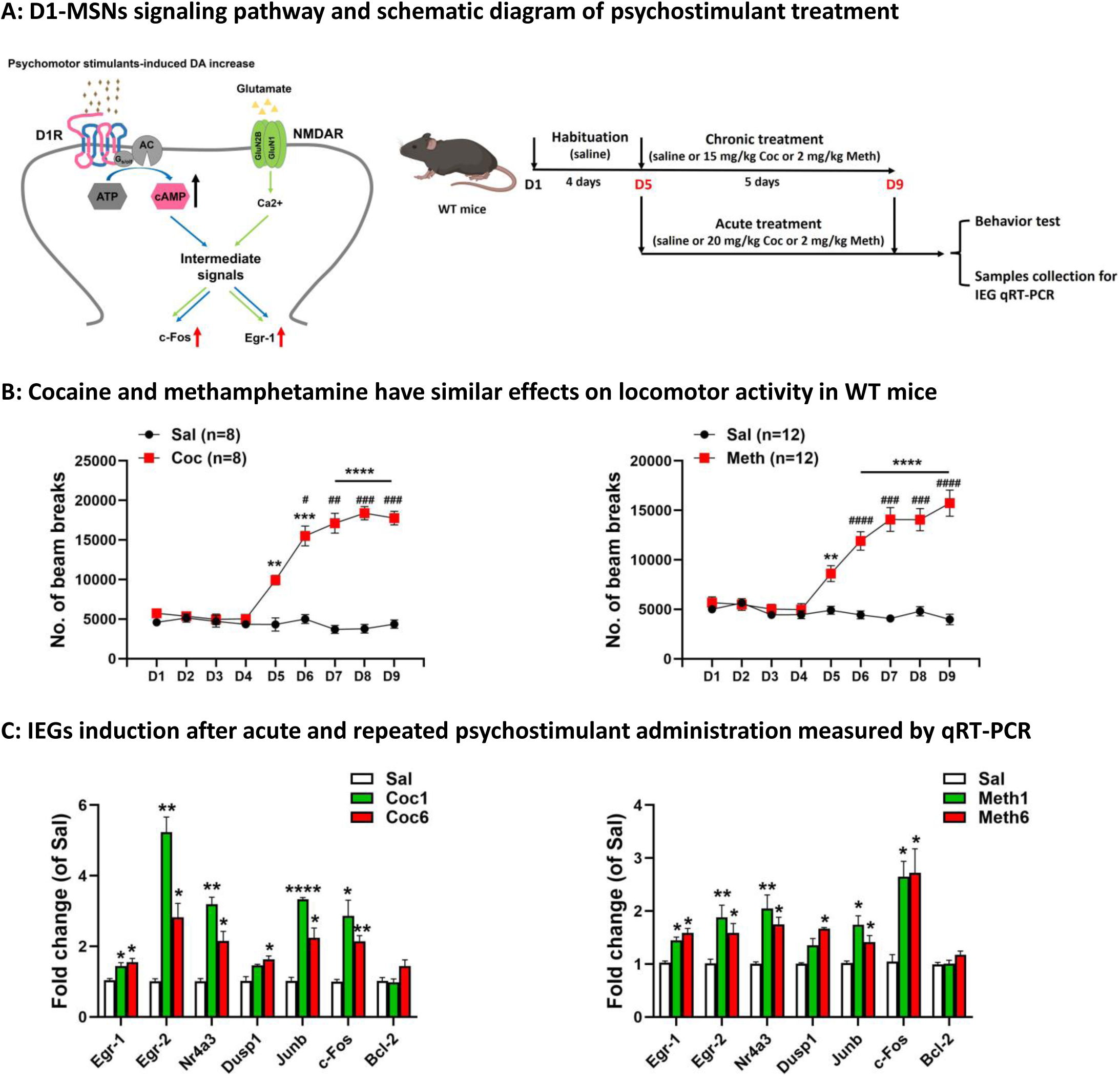
Cyclic AMP dependence of IEG induction after psychostimulant administration. (A) Schematic model of D1 receptor→cyclic AMP (cAMP) metabotropic and glutamate→ calcium inotropic signaling pathway mediated immediate early genes induction in D1 medium spiny neurons (MSNs) after psychostimulant administration (Left) and acute and repeated cocaine (Coc, 20 or 15 mg/kg, i.p) or methamphetamine (Meth, 2 mg/kg, i.p) treatment for 5 days in WT mice for the measurement of locomotor activity and IEG expression by qRT-PCR (Right). (B) Mice developed robust locomotion sensitization both by cocaine (Coc, Left) and methamphetamine (Meth, Right) for 5 consecutive injections. n = 8 mice in Coc group and n = 12 mice in Meth group, **p < 0.01, ***p < 0.001, ****p < 0.0001 for Coc or Meth versus saline at each time point. #p < 0.05, ##p < 0.01, ###p < 0.001, ####p < 0.0001 for Coc or Meth D6-D9 versus D5. All panels: Mean ± S.E.D. On day 9 (D9) after 1 hr behavior recording, mice were sacrificed and NAc were dissected for qRT-PCR assay (shown in Supple Figure 1C) (C) Induction of IEG by acute (Coc1 or Meth1) or repeated cocaine or methamphetamine (Coc6 or Meth6) measured by qRT-PCR. n = 5∼8 mice in each group, *p < 0.05, **p < 0.01, ****p < 0.0001 for Coc or Meth versus saline. All panels: Mean ± S.E.D.

**Supplementary Figure 2.**
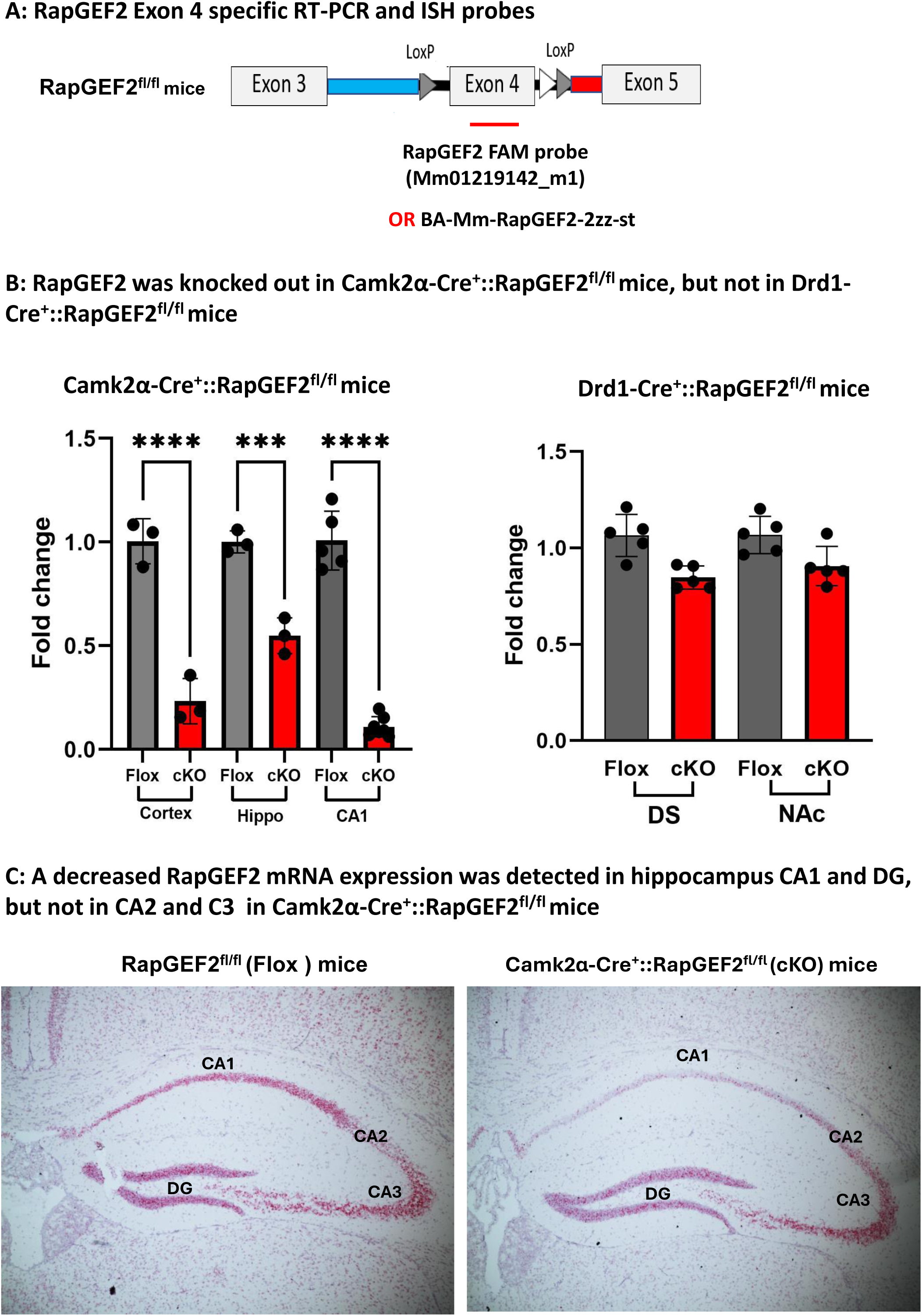
Camk2a-Cre, but not Drd1-Cre, can penetrate LoxP locus in RapGEF2 flox mice to knock-out RapGEF2. (A) Design of the RapGEF2 Exon 4 specific TaqMan probe for qRT-PCR and Basescope probe for ISH, which allow us to check the RapGEF2 expression difference between control flox mice and Cre::RapGEF2^fl/fl^ mice. (B) In Camk2α-Cre::RapGEF ^fl/fl^ mice, the relative level of RapGEF2 mRNA was ∼70% decrease in the cortex, ∼50% decrease in the whole hippocampus and ∼85% decrease in hippocampus CA1, compared to Flox control mice. N = 5∼8 mice for each group. ***p < 0.001, ***p < 0.0001 for cKO versus Flox mice. All panels: Mean ± S.E.D. Similarly, detection of RapGEF2 mRNA (red color) in hippocampus showed a significant decrease of RapGEF2 mRNA in CA1 and DG, but not in CA2 and CA3, indicating the specificity of this custom-designed RapGEF2 Basescope prpbe. which is consistent with previous data shown by Western blot and immunohistochemistry (Jiang SZ JN 2021). (C) However, in the Drd1-Cre::RapGEF ^fl/fl^ mice, only a decrease tread of RapGEF mRNA was seen in the cKO by qRT-PCR (N = 5 mice for each group). Further ISH data demonstrated RapGEF2 mRNA was only knock out in some of the D1-neurons (shown by open arrows) and most of the D1-neurons still had RapGEF2 expression (shown by solid arrows), indicating the less efficiency of Drd1-Cre to penetrate the LoxP locus of RapGEF2 flox mice to knock out the RapGEF2.

**Supplementary Figure 3.**
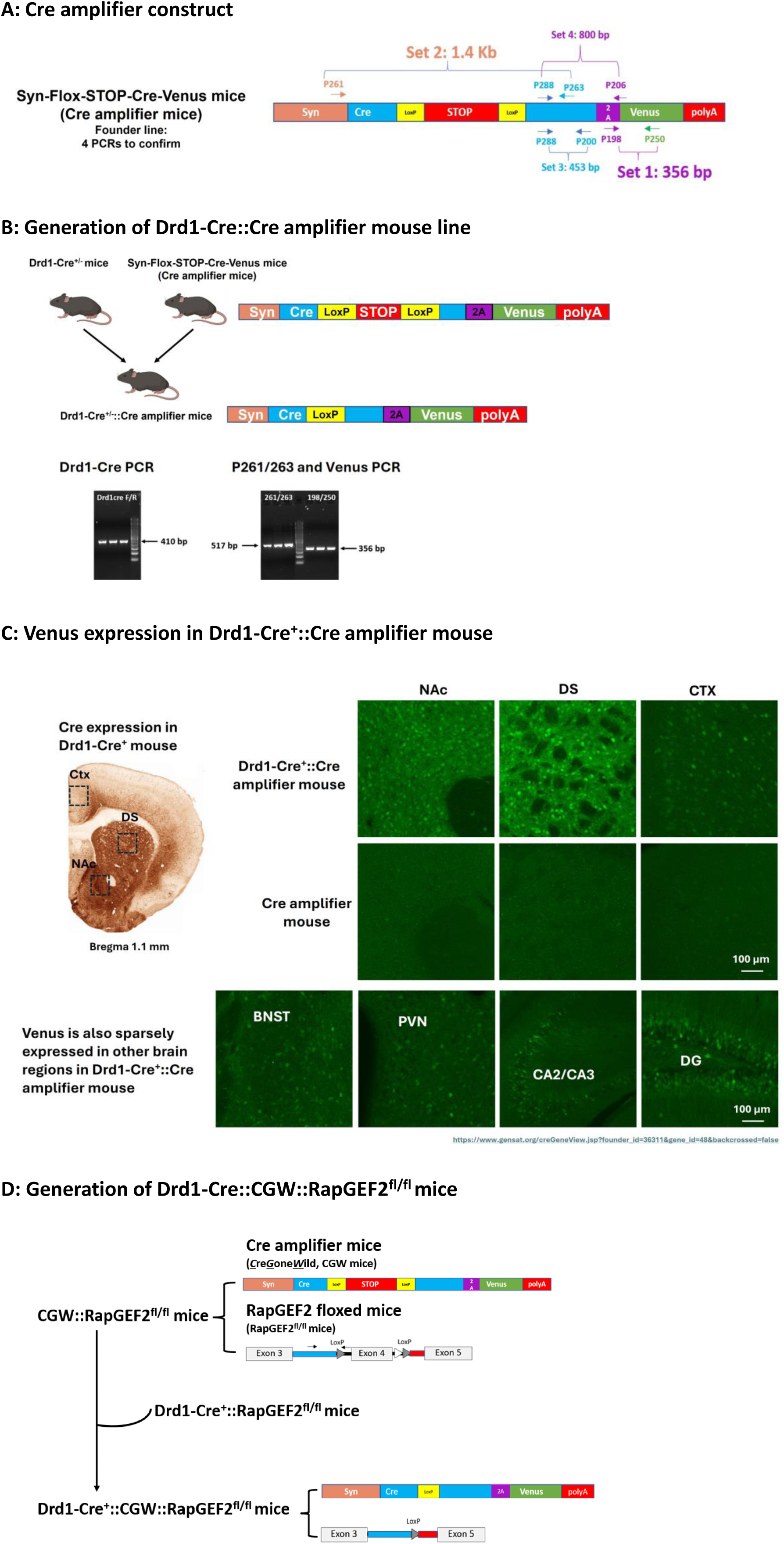
Generation of Cre amplifier mouse and D1-specific RapGEF2 knock-out mouse. (A) Cre dependent Cre amplifier construct. A “floxed” STOP cassette, flanked by two loxP sites, was inserted into Cre sequence driven by a Synapsin promoter to generate a Cre dependent Cre amplifier mouse, also call Syn-Flox-STOP-Cre-Venus construct, which have Venus expression with presence of Cre. (B) Generation of and Drd1-Cre::Cre amplifier mouse by crossing Drd1-Cre^+^ mouse with Cre amplifier mouse, in which the STOP cassette are excised to allow more Cre expression and light up the D1-MSNs with Venus expression. Two PCR are required to genotype the offspring to detect the presence of both Drd1-Cre and Syn-Flox-STOP-Cre. (C) Venus expression was only detected in Drd1-Cre^+^::Cre amplifier mouse, but not Cre amplifier only mouse, by using a GFP antibody in multiple brain area, including very dense expression in dorsal and ventral striatum and sparsely low expression in prefrontal cortex, the bed nucleus of the stria terminalis (BNST), the paraventricular nucleus (PVN), hippocampus CA2/CA3 and dentate gyrus (DG) area, which is consistent with the Cre expression pattern in Drd1-Cre mouse (referred to GENSAT Drd1-Cre FK150). (D) Generation of Drd1-Cre::Cre amplifier::RapGEF2^fl/fl^ mice by breeding Cre amplifier (also called CreGoneWild, CGW) or Drd1-Cre with RapGEF2^fl/fl^ mice first to generate CGW::RapGEF2^fl/fl^ and Drd1-Cre::RapGEF^fl/fl^ mice, respectively, then breed them together to knock-out RapGEF2 Exon4 specifically in D1 neurons.

**Supplementary Figure 4.**
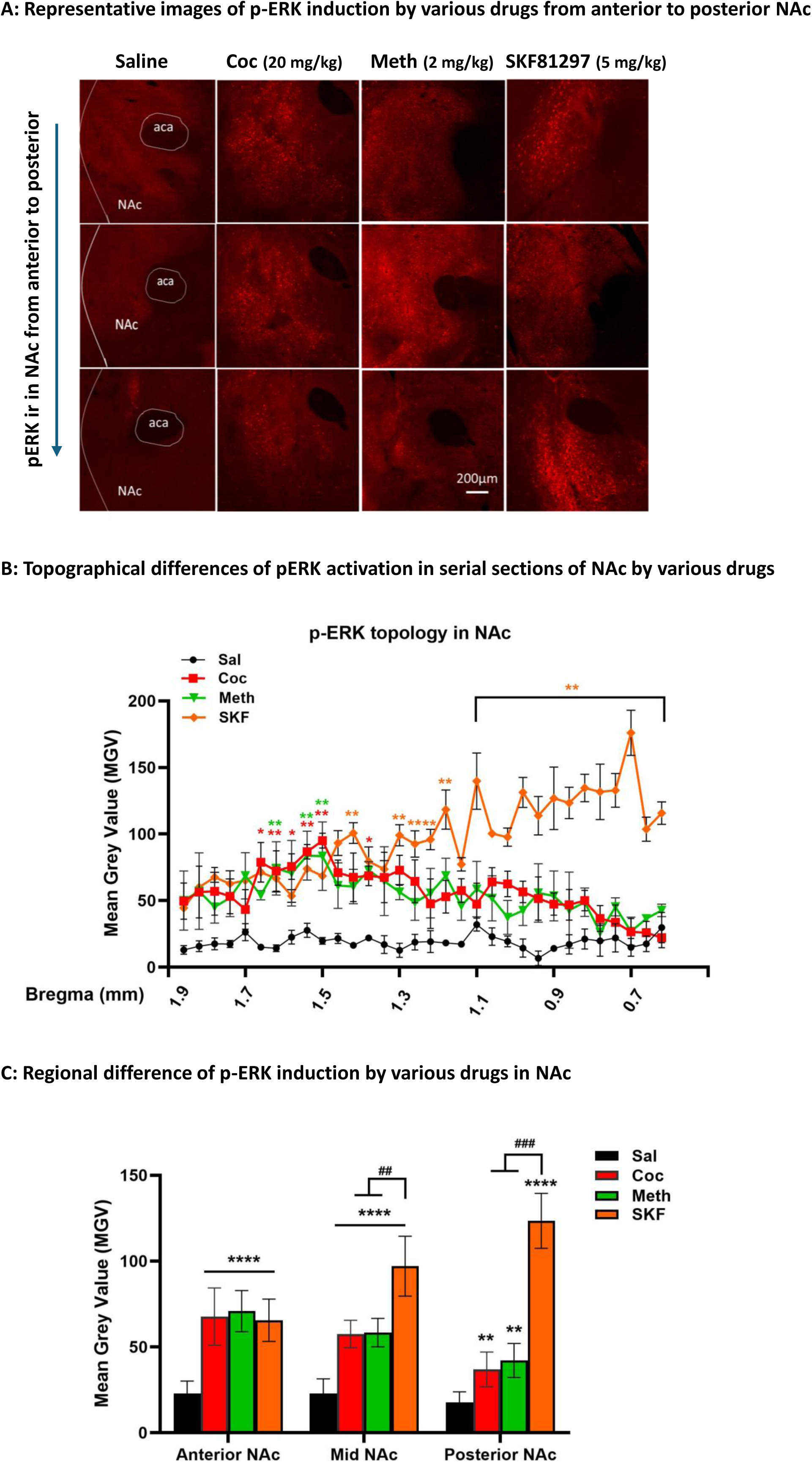
p-ERK induction pattern by psychostimulants and D1-agonist in NAc. (A) Representative images of p-ERK induction by single dose of cocaine (20 mg/kg, Coc), methamphetamine (2 mg/kg, Meth) and D1-agonist SKF81297 (SKF, 5 mg/kg) from anterior to posterior NAc at 10∼15 min after drug injection (i.p), demonstrating that induction of p-ERK was mainly in anterior and medial part of NAc for cocaine and methamphetamine and medial and posterior part of NAc for SKF81297. (B) Topographical differences of pERK activation by Coc, Meth and SKF in NAc from Bregma 1.9 mm to 0.7 mm, a total of 32 sections (40 um for each), showing that a significant increase of p-ERK expression was found in NAc from 1.5 mm to 1.2 mm by 3 drug treatment and only SKF81297 induced more p-ERK expression in the NAc from 1.2 mm to 0.7 mm, compared to Coc or Meth and Sal. N=3 mice in each group. *p < 0.05, **p < 0.01 for Coc or Meth or SKF versus Sal. All panels: Mean ± S.E.D. (C) Reginal differences of pERK induction by Coc, Meth and SKF in NAc, demonstrating that all 3-drug treatment increased pERK expression throughout NAc and SKF81297 induced more p-ERK expression in medial and posterior NAc, compared to Coc or Meth. N=3 mice in each group. **p < 0.01, ****p < 0.0001 for Coc or Meth or SKF versus Sal. ##p < 0.01, ####p < 0.0001 for SKF versus Coc or Meth. All panels: Mean ± S.E.D.

